# PlasClass improves plasmid sequence classification

**DOI:** 10.1101/783571

**Authors:** David Pellow, Itzik Mizrahi, Ron Shamir

## Abstract

**Background:** Many bacteria contain plasmids, but separating between contigs that originate on the plasmid and those that are part of the bacterial genome can be difficult. This is especially true in metagenomic assembly, which yields many contigs of unknown origin. Existing tools for classifying sequences of plasmid origin give less reliable results for shorter sequences, are trained using a fraction of the known plasmids, and can be difficult to use in practice.

**Results:** We present PlasClass, a new plasmid classifier. It uses a set of standard classifiers trained on the most current set of known plasmid sequences for different sequence lengths. PlasClass outperforms the state-of-the-art plasmid classification tool on shorter sequences, which constitute the majority of assembly contigs, while using less time and memory.

**Conclusions:** PlasClass can be used to easily classify plasmid and bacterial genome sequences in metagenomic or isolate assemblies. It is available from: https://github.com/Shamir-Lab/PlasClass

## Background

When using high-throughput sequencing to study the presence and dynamics of plasmids in their bacterial hosts, it is often necessary to classify sequences as being of plasmid or bacterial origin. This is especially true in the case of metagenomic sequencing, which can include many sequences of unknown origin and varying lengths. We focus on the challenge of classifying contigs in a metagenomic assembly in order to identify which are of plasmid origin.

The current state-of-the-art classifier of plasmid sequences is PlasFlow [1], a neural network based algorithm that was shown to perform better than previous tools such as cBar [2]. While PlasFlow is successful in classifying small sets of long sequences, it produces less reliable results for short sequences and requires large memory on very large metagenomic datasets.

Here we present PlasClass, a new plasmid sequence classifier implemented as an easy to use Python package. It uses a set of logistic regression classifiers each trained on sequences of a different length sampled from plasmid and bacterial reference sequences. When applied on a set of sequences, the appropriate length-specific classifier is used for each sequence.

We tested PlasClass on simulated data, on bacterial isolates, on a wastewater plasmidome, and on plasmids assembled from human gut microbiome samples. For shorter sequences, which are the majority of contigs in an assembly, PlasClass achieved better F1 scores than PlasFlow. It used significantly less RAM and disk memory, and can be run much faster by using multiprocessing.

PlasClass is provided at https://github.com/Shamir-Lab/PlasClass.

## Implementation

### Training databases

We used reference sequence databases to obtain the training sequences for our classifiers. For the plasmid references we used plasmid sequences listed in PLSDB [3] (v.2018_12_05), an up-to-date curated plasmid database. After filtering out duplicate sequences this database contained 13469 reference plasmids (median length: 53.8kb).

For the bacterial references we downloaded all complete bacterial genome assemblies from NCBI (download date January 9, 2019). We removed sequences annotated as being plasmids and filtered out duplicates, leaving 13491 reference chromosomes (median length: 3.7Mbp).

One quarter of the sequences were randomly removed from the databases before training in order to provide a held-out test set for validation. PlasClass was retrained on the full databases and this version was used for testing on assembled data.

### Training the classifiers

We sampled sequence fragments of different lengths from the reference sequences with replacement and constructed a k-mer frequency vector for each fragment. Canonical k-mers of lengths 3-7 were used, resulting in a feature vector of length 10952 for each fragment. Fragment lengths were 500k, 100k, 10k, and 1k. For the two shorter lengths, 90,000 training fragments were used from each class. For the lengths 500k and 100k, since there were not enough long plasmids to do the same, we sampled enough fragments to cover all of the sufficiently long plasmids to a depth of 5. This resulted in 1934 and 45525 plasmid fragments of length 500k and 100k, respectively on the full plasmid database.

For each length, a logistic regression classifier was trained on the plasmid and bacterial fragments’ k-mer frequency vectors using the scikit-learn [4] machine learning library in Python. Code is provided to retrain the models on user-supplied reference sequence databases.

### Length-specific classification

PlasClass uses four logistic regression models to classify sequences of different length. Each sequence is assigned to the closest length from among 1kb, 10kb, 100kb, and 500kb. Equivalently, this defines four length ranges: (0,5.5kb], (5.5kb,55kb], (55kb,300kb], (300kb, ∞). Given a sequence, its k-mers are counted, the canonical k-mer frequency vector is calculated and used to classify it with the classifier for the range it falls into. k-mer counting can be performed in parallel for different sequences. Finally, all classification results are concatenated into a single output in the same order as the input sequences.

### Classification with PlasClass

PlasClass is available at https://github.com/Shamir-Lab/PlasClass. It has been retrained using the full set of database references. PlasClass can be used as a command-line tool to classify sequences in an input fasta file or it can be imported as a module into the user’s code to classify sequences in the user’s program. It can be run in parallel mode to achieve faster runtimes. PlasClass is fully documented in the Supplement (Section S3) and at the url provided above.

## Results

We compared performance of PlasFlow and PlasClass on both simulated and real data.

### Experimental settings

PlasFlow and PlasClass both assign class probabilities to each sequence. We say a sequence is classified as having plasmid origin if the probability that it belongs to the plasmid class is > 0.5. When running PlasFlow, this probability was summed over all plasmid classes, and we set the parameter `--threshold=0.5` to ensure each sequence is classified as either plasmid or bacterial. All assemblies were performed using the `--meta` option of SPAdes [5] v3.12.

### Performance metrics

We calculated the precision, recall and F1 scores counting the *number* of true positive and false positive predictions. Some previous works ([6, 1]) calculated performance based on the *lengths* of the sequences classified as plasmids and the total length of the plasmids in a sample. A length-weighted metric is appropriate in the context of plasmid sequence assembly, but in the context of contig classification this makes little sense. (Consider the extreme case of one extremely long sequence and 999 very short ones. Classifying the long contig is easy, but a classifier that only identifies it correctly will have weighted precision and recall near 1 even though only 1/1000 of the sequences are correctly classified.) For this reason we used the numbers of correctly classified sequences.

Our prediction problem is particularly challenging for two reasons. The first is that when classifying contigs produced in assembly, most contigs are very short and thus hard to classify. The second is the acute class imbalance between the number of plasmid and bacterial contigs. Bacteria are roughly two orders of magnitude larger than plasmids, as was the case in our training databases. As a result, the fraction of plasmid contigs in isolate and metagenomic classification setups is also tiny. Consider an assembly with 10,000 contigs, of which 100 are of plasmid origin. A random baseline, which predicts that a sequence is of plasmid origin with 50% probability, will correctly classify 50 of the plasmid contigs on average, achieving a recall of 50%. However, it will also classify on average half of the bacterial contigs as plasmids, achieving a precision of only 1% and an F1 score of 2%. An alternative random baseline using the correct 1% probability for plasmid sequences would achieve precision, recall and F1 of 1%. So one would expect substantially lower performance numbers than in balanced classification setups.

### Classifying sequences from held-out references

We sampled overlapping L-long fragments covering the held out plasmids with an overlap of *L*/2 for *L* = 100k, 10k and 1k. A matching number of *L*-long fragments were sampled from the held out bacterial genomes for each length *L*. (Note that this creates a balanced classification scenario.) Table 1 summarizes the classification results. PlasClass improved precision at the cost of slightly lower recall and had better overall F1 on the shorter sequence lengths. These short sequences can make up the majority of contigs in metagenomic assemblies (see Tables 2, 4 and Figure S1), allowing PlasClass to outperform PlasFlow in many settings as shown below.

**Table 1:** Performance of PlasFlow and PlasClass on fixed length sequence fragments sampled from the held out references.

| Length (bp) | # fragments<br>per class | PlasFlow |  |  | PlasClass |  |  |
| --- | --- | --- | --- | --- | --- | --- | --- |
|  |  | Precision | Recall | F1 | Precision | Recall | F1 |
| 100k | 2979 | 95.6 | 88.4 | 91.9 | 96.9 | 85.4 | 90.8 |
| 10k | 56583 | 83.1 | 87.7 | 85.3 | 88.7 | 86.4 | 87.6 |
| 1k | 607656 | 59.7 | 79.1 | 68.1 | 75.1 | 74.6 | 74.8 |

### Performance on a benchmark of bacterial isolates

We compared the performance of PlasFlow and PlasClass on the isolate assemblies from the benchmark in [6]. Specifically, we downloaded the assemblies and all bacterial and plasmid reference sequences used in the benchmarking experiment of [6] (available from: https://gitlab.com/sirarredondo/Plasmid_Assembly). Assembled contigs were mapped to the references using BLAST and only contigs that uniquely mapped to either the plasmid or chromosome class were retained (>95% mapping identity along >95% of the contig length). There were 60579 contigs across all the assemblies of which 27677 uniquely mapped to only one class (74 plasmid and 27603 chromosome) and were used in this test. As seen in Table 2, the majority of these sequences were extremely short (66% of the 27677 contigs <500bp). In this hard classification challenge, both classifiers obtained very low precision, but PlasClass had consistently higher F1. Notably, the results of both methods improved when shorter sequences were filtered out. In S2 we demonstrate the effect of alternative performance metrics, which result in much higher scores.

**Table 2:** Performance on bacterial isolates from [6], as a function of the minimum contig length.

| Contig length (bp) | # of contigs | PlasFlow |  |  | PlasClass |  |  |
| --- | --- | --- | --- | --- | --- | --- | --- |
|  |  | Precision | Recall | F1 | Precision | Recall | F1 |
| All | 27677 | 0.401 | 90.54 | 0.798 | 0.751 | 87.84 | 1.49 |
| >500 | 9292 | 1.17 | 91.49 | 2.31 | 2.26 | 91.74 | 4.41 |
| >1000 | 5762 | 1.78 | 88.57 | 3.49 | 3.09 | 94.29 | 5.98 |
| >5000 | 3481 | 2.72 | 93.33 | 5.29 | 4.72 | 100 | 9.01 |

### Performance on simulated metagenome assemblies

We simulated metagenomes by randomly selecting bacterial genome references from the NCBI along with their associated plasmids and using realistic distributions for genome abundance and plasmid copy number. For genome abundance we used the log-normal distribution, normalized so that the relative abundances sum to 1. For plasmid copy number we used a geometric distribution with parameter *p* = *min*(1, *log*(*L*)/7) where *L* is the plasmid length. This makes it much less likely for a long plasmid to have a copy number above 1, while shorter plasmids can have higher copy numbers. Short reads were simulated from the genome references using InSilicoSeq [7] and assembled.

We then classified the assembled contigs. Classification was performed on the assembled contigs that had a unique and unambiguous match to either a reference plasmid or reference chromosome sequence used in the simulation (1120 plasmid contigs, 32451 chromosome contigs in Sim1, and 11579 plasmid contigs, 374397 chromosome contigs in Sim2). F1 results are shown in Table 3. PlasClass outperformed PlasFlow by more than 25%. Scores were low for both methods due to the many short contigs in the assembly (49% and 72% of the contigs <500 bp in Sim1 and Sim2 respectively) and the class imbalance. We show the impact of short sequences on performance in Table 4. PlasClass consistently outperformed PlasFlow, and both methods performed better as more short sequences were filtered out.

**Table 3:** Summary of the simulated metagenome datasets and comparison of F1 scores. # unique is the number of distinct plasmids, ignoring multiple copies.

|  | # chromosomes | # plasmids | # unique | # contigs | PlasFlow F1 | PlasClass F1 |
| --- | --- | --- | --- | --- | --- | --- |
| Sim1 | 34 | 82 | 56 | 33571 | 9.67 | 12.32 |
| Sim2 | 198 | 333 | 219 | 385976 | 7.31 | 10.79 |

**Table 4:** Performance on simulated metagenomes as a function of the minimum contig length.

|  | Contig length (bp) | # of contigs | PlasFlow |  |  | PlasClass |  |  |
| --- | --- | --- | --- | --- | --- | --- | --- | --- |
|  |  |  | Precision | Recall | F1 | Precision | Recall | F1 |
| Sim1 | All | 33571 | 5.13 | 88.21 | 9.69 | 6.73 | 72.59 | 12.32 |
|  | >500 | 16973 | 7.61 | 86.27 | 13.99 | 10.64 | 80.07 | 18.79 |
|  | >1000 | 11665 | 10.36 | 86.30 | 18.50 | 14.98 | 81.60 | 25.31 |
|  | >5000 | 4029 | 26.79 | 89.47 | 41.23 | 35.79 | 86.64 | 50.65 |
| Sim2 | All | 385976 | 3.82 | 84.93 | 7.31 | 5.83 | 71.97 | 10.79 |
|  | >500 | 106388 | 8.02 | 86.21 | 14.67 | 13.35 | 79.25 | 22.85 |
|  | >1000 | 45323 | 13.41 | 89.15 | 23.31 | 21.93 | 83.58 | 34.75 |
|  | >5000 | 5579 | 38.48 | 94.68 | 54.72 | 46.39 | 87.41 | 60.61 |

### Performance on plasmidome sample

We assembled the wastewater plasmidome sample ERR1538272 from the study by Shi et al. [8]. It is a metagenomic sample that was enriched for plasmid sequences. Each contig in the assembly was matched to the plasmid and bacterial reference databases using BLAST. The set of 9118 contigs (out of 35285) that uniquely matched only one of the databases (1152 plasmid contigs, 7966 chromosome contigs) was used as the gold standard to test the classifiers (contig length distribution is presented in the Supplement, Figure S1). Although the plasmid-enriched setting favors PlasFlow, which sacrifices precision for higher recall, PlasClass still had a higher combined F1 as shown in Table 5.

**Table 5:**
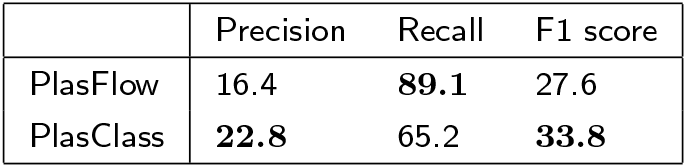
Performance of PlasClass and PlasFlow on the plasmidome sample

|  | Precision | Recall | F1 score |
| --- | --- | --- | --- |
| PlasFlow | 16.4 | **89.1** | 27.6 |
| PlasClass | **22.8** | 65.2 | **33.8** |

We computed the precision-recall curve for the classification of the gold standard contigs in this sample by PlasClass, shown in Figure S2. The area under the curve is 0.29, more than double the baseline of 0.13 (the fraction of the contigs that are of plasmid origin).

### Classifying plasmids assembled from metagnomic samples

We assembled six publicly available human gut microbiome samples (accessions: ERR1297700, ERR1297720, ERR1297770, ERR1297796, ERR1297822, ERR1297834) and found plasmid sequences in the assemblies using Recycler [9]. Recycler assembles plasmid sequences based on coverage and circularity - features that are not used by the classifiers. 16-27 plasmids were assembled per sample (median length: 3.4kb). We classified each of the plasmids generated by Recycler to determine the extent of agreement between the sequence classifiers and this orthogonal approach. As seen in Figure 1, PlasClass agreed with Recycler on the same number or more plasmids than PlasFlow in all samples. This suggests that PlasClass can correctly identify more plasmids in real datasets, which contain many previously unknown plasmid sequences.

**Figure 1:**
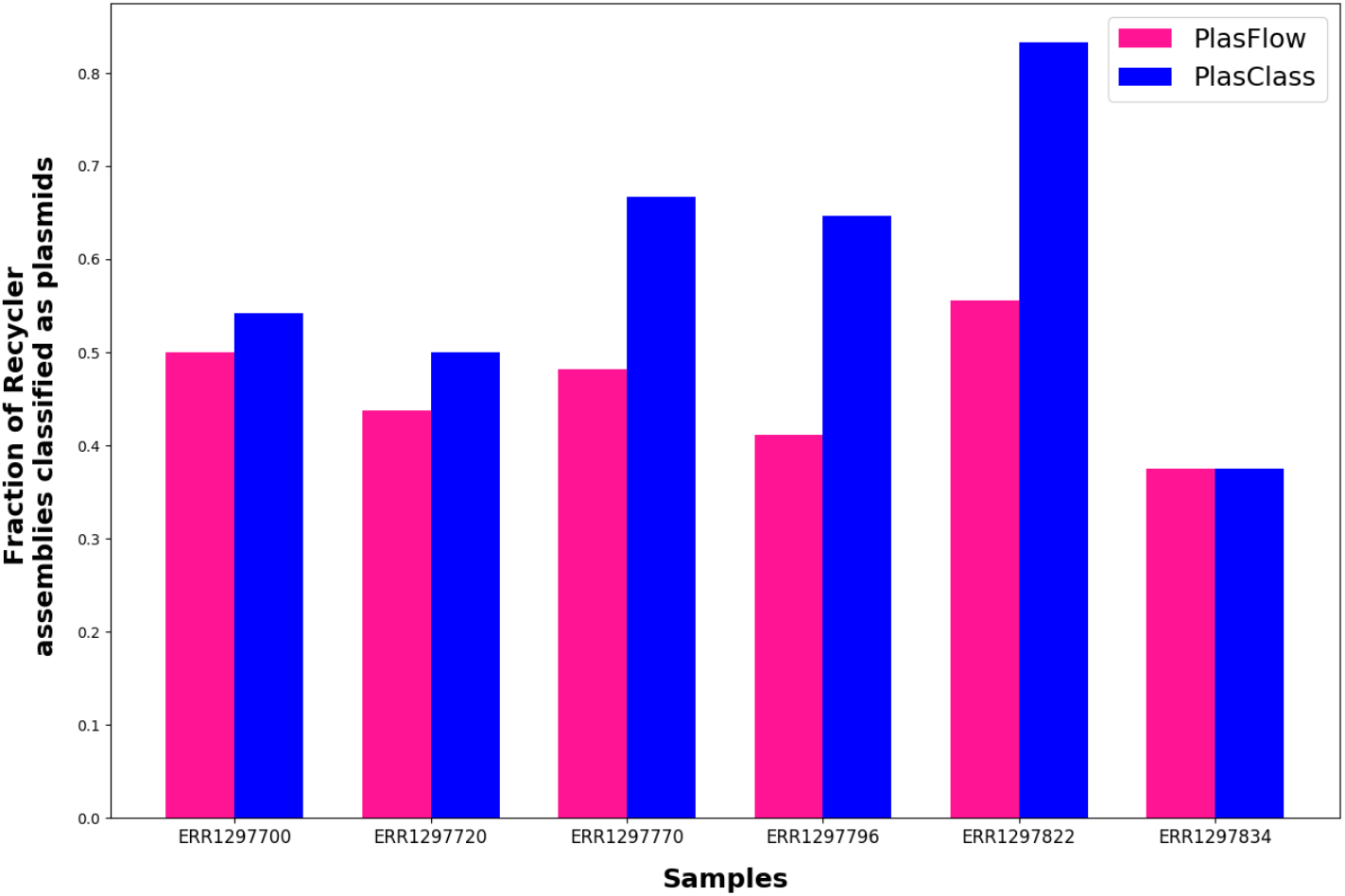
Agreement of PlasFlow and PlasClass classifications with the plasmids generated by Recycler.

### Resource usage

In Table 6, we compare the runtime and memory usage of PlasClass and PlasFlow on the full plasmidome, simulated metagenome, and isolate bacterial datasets. PlasClass (running with a single process) was faster than PlasFlow on the most time consuming sample and was significantly faster in all cases when using multipro-cessing. It used less than half the RAM of PlasFlow and the RAM usage was not increased significantly when using multiprocessing. PlasFlow writes the feature matrices to disk while PlasClass does not. Performance was measured on a 44-core, 2.2 GHz server with 792 GB of RAM.

**Table 6:** Runtime (wall clock time, in minutes) and memory usage (in GB) of PlasClass and PlasFlow.

| Dataset | PlasFlow |  |  | PlasClass |  | PlasClass - 8 processes |  |
| --- | --- | --- | --- | --- | --- | --- | --- |
|  | Runtime | RAM | Disk | Runtime | RAM | Runtime | RAM |
| Isolates | 12.8 | 47.8 | 21.4 | 36.3 | 17.2 | 6.8 | 17.2 |
| Sim1 | 7.1 | 28.3 | 12.1 | 16.2 | 12.0 | 3.0 | 12.0 |
| Sim2 | 89.3 | 291.3 | 137.5 | 54.8 | 17.3 | 17.1 | 17.3 |
| Plasmidome | 7.9 | 28.8 | 12.2 | 4.2 | 12.2 | 5.2 | 17.3 |

### Summary and discussion

We presented the PlasClass classifier and have shown that it outperforms the state-of-the-art classifier in identifying plasmid sequences. Classifying plasmid sequences in the real-world context of metagenomic data is a difficult task due to the nature of the assembled sequences: the sequences are mostly short (60-90% are shorter than 1 kbp, see Tables 2 and 4), and there is acute imbalance between the number of plasmid and bacterial sequences (1:500 in the bacterial isolates, and 1:8 even in the plasmid-enriched plasmidome sample). We showed that in cases where short sequences are filtered out or when plasmid sequences are enriched, sequence classifiers are able to achieve good performance.

Although performance scores are low, we showed that PlasClass performs relatively better than PlasFlow in most cases. The low performance scores are also due in part to our careful construction of the gold standard datasets used for testing. In Supplement S2, we demonstrated that when we instead consider all sequences that could be of either plasmid or bacterial origin to belong to the plasmid class, as done in [1], the scores are much higher. In this case, we achieved F1 scores in the range of 25-85%. As in the case with the more strict metrics, PlasClass performed better than PlasFlow, and by a similar factor.

Overall, PlasClass’s classification performance is better than or similar to that of PlasFlow across a wide range of contexts and it demonstrates much faster runtimes with significantly lower memory usage.

## Declarations

### Authors’ contributions

DP and RS conceived this study. DP implemented the classifier, and performed the analysis. RS and IM supervised the analysis. All authors wrote the manuscript.

## Acknowledgements

We thank the members of the Shamir and Mizrahi Labs for their help and advice.

## Additional Files

Additional file 1 — Supplementary Information.pdf

Supplementary information for: PlasClass improves plasmid sequence classification

