## Supplementary Information.pdf for "PlasClass improves plasmid sequence classification"

### Supplementary information for:
### PlasClass improves plasmid sequence classification

#### S1 Plasmidome sample

Figure S1 presents the distribution of contig lengths in the assembly graph of the wastewater plasmidome sample ERR1538272 from the study by Shi et al. [1].

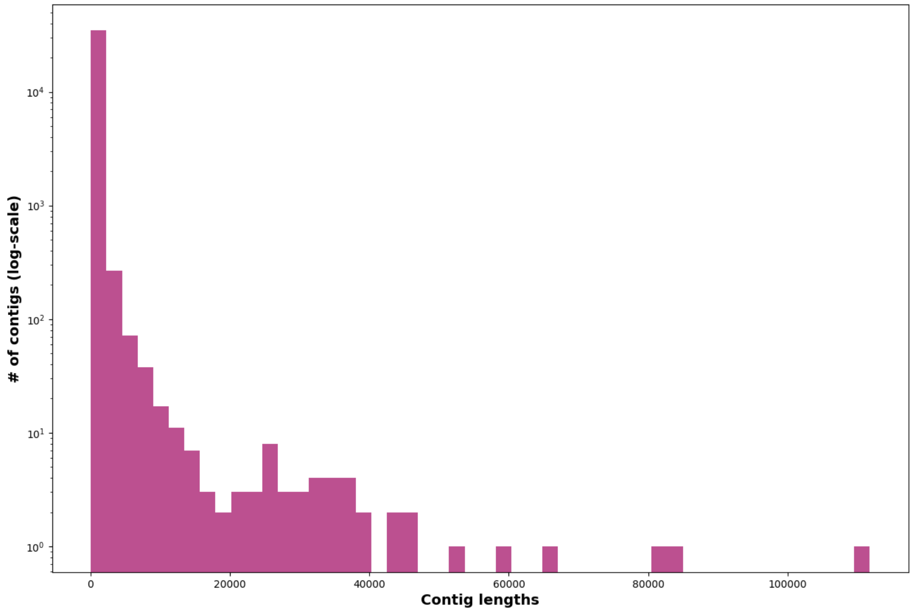


Figure S1: Histogram of the contig lengths in the plasmidome assembly. Note that the y-axis uses log-scale.

Figure S2 presents the precision-recall curve for the classification of the plasmidome sample by PlasClass. The area under the precision-recall curve is AUPR=0.29. The baseline given the class imbalance is 0.13.

#### S2 Alternative performance metrics

The performance numbers of both PlasFlow and PlasClass were low across most of the datasets tested, especially in the isolates data. The reasons for this were explained in the section on performance metrics. The numbers reported by PlasFlow on this data were significantly higher due to a number of factors, including two major ones: short sequences were filtered out, and contigs that matched both plasmid and bacteria references were considered to be plasmids.

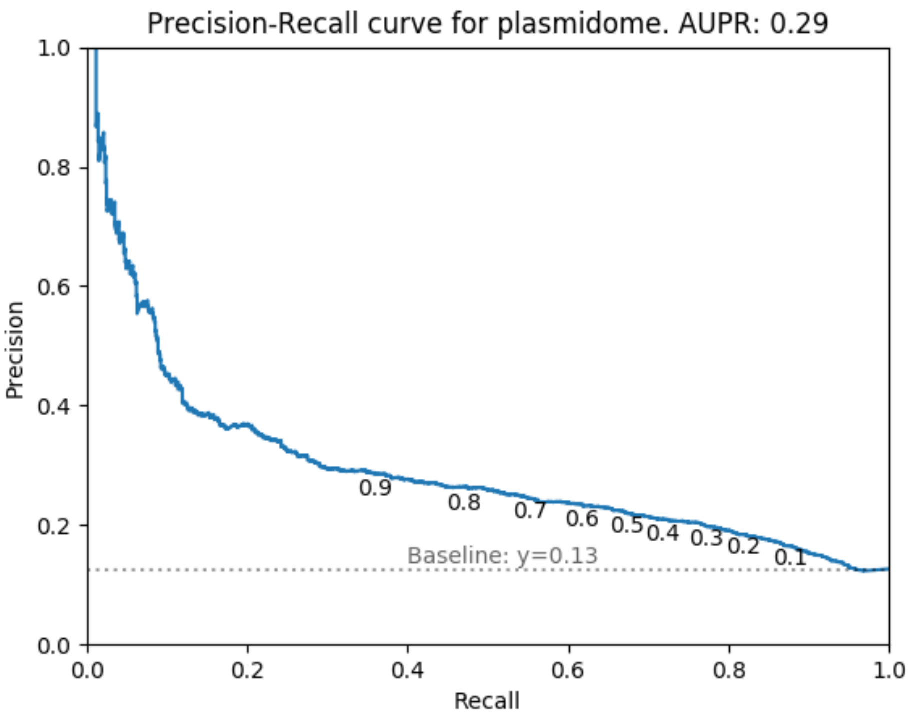


Figure S2: Precision-recall curve for the classification of contigs of unambiguous plasmid or chromosome origin in the plasmidome sample.

Considering contigs matching both classes as plasmids could be appropriate for classifying all sequences in an assembly to determine their origin. However, in using these assemblies to create a gold-standard dataset for benchmarking a classifier, it is more suitable to filter out ambiguous sequences that may belong to both classes.

Table S1 reports the performance on the bacterial isolates dataset when contigs matching both classes are considered as plasmids, and when filtering out short sequences.

Table S1: Performance calculated using the alternative procedure on the isolate bacteria dataset. Here, contigs matching both classes were considered plasmids, and sequences shorter than 1kb were excluded.

| | PlasFlow | | | PlasClass | | |
|---|---|---|---|---|---|---|
| Classified data | Precision | Recall | F1 | Precision | Recall | F1 |
| Only unambiguous matches | 0.401 | 90.54 | 0.798 | 0.751 | 87.84 | 1.49 |
| All plasmid matches are positive | 31.16 | 87.77 | 46.0 | 43.65 | 77.58 | 55.87 |
| Plasmid matches positive, $>1$ kbp | 47.54 | 90.04 | 62.22 | 51.95 | 91.82 | 72.54 |

When using this alternative performance yardstick, we observed much higher values, consistent with previously reported results. The same was true on the other datasets as well. Table S2 reports performance on the simulated metagenomes and plasmidome when using this alternative calculation.

Table S2: Performance calculated using the alternative procedure on simulated metagenomes and plasmidomes. All plasmid matches were considered to be positive examples and contigs $< 1$ kbp were filtered out.

| | PlasFlow | | | PlasClass | | |
|---|---|---|---|---|---|---|
| Sample | Precision | Recall | F1 | Precision | Recall | F1 |
| Sim1 | 10.9 | 85.0 | 19.4 | 15.7 | 81.0 | 26.3 |
| Sim2 | 14.0 | 86.5 | 24.1 | 22.4 | 79.2 | 35.0 |
| Plasmidome | 78.8 | 86.8 | 82.6 | 87.0 | 81.5 | 84.2 |

Using this alternative measure, PlasClass consistently outperformed PlasFlow, as we observed with the regular performance measure, and by a similar factor.

#### S3 PlasClass documentation

PlasClass is fully documented at `https://github.com/Shamir-Lab/PlasClass`. We outline the installation and usage instructions here.

##### Installing PlasClass

PlasClass is written in Python3 and requires NumPy and scikit-learn. To install PlasClass do:

```
git clone https://github.com/Shamir-Lab/PlasClass.git
cd PlasClass
python setup.py install
```

We recommend using a virtual environment.

##### Classifying sequences in a fasta file

The `classify_fasta.py` script can be used to classify the sequences in a fasta file. It is invoked as follows:

```
python classify_fasta.py -f <input file> [-o <output file>] [-p <# processes>]
```

The required option is:

- `-f/--fasta`: String - the fasta file to be classified.

The two optional options are:

- `-o/--outfile`: String - the name of the output file.
  Default: `<in>.probs.out` (`<in>` is the path to the fasta file).
- `-p/--num_processes`: Integer - the number of processes to use. Default: 8.

The output file is a tab separated file with each line containing a sequence header and the corresponding score. The sequences are in the same order as in the input fasta file.

##### Calling the PlasClass module

The classifier can also be imported and called directly in a Python program. For example, once the `plasclass` module has been installed the following lines of code can be used:

```
from plasclass import plasclass
my_classifier = plasclass()
my_classifier.classify(seqs)
```

The `plasclass()` constructor takes 3 optional parameters:

- `n_procs`: Integer - Number of processes to use (default: `1`)
- `scales`: Integer array - Scales of the sequence length bins (default: `[1000,10000,100000,500000]`)
- `ks`: Integer array - Values of the k-mer lengths to use (default: `[3,4,5,6,7]`)

The sequence(s) to classify, `seqs`, can be either a single string or a list of strings. The strings must be uppercase. The function `plasclass.classify(seqs)` returns a list of plasmid scores, one per input sequence, in the same order as the input.

##### Training PlasClass models

The user can train models by providing two fasta files with plasmid and bacterial reference sequences to the `train.py` script. Required options are:

- `-p/--plasmid`: String - the fasta file of the plasmid references.
- `-c/--chromosome`: String - the fasta file of the chromosome references.
- `-o/--outdir`: String - the path of the output directory.
  Default: directory of script.

The optional options are:

- `-n/--num_processes`: Integer - the number of processes to use.
  Default: 16.

- `-k/--kmers`: String - comma separated list of the k-mer sizes to use.
  Default: 3,4,5,6,7.
- `-l/--lengths`: String - comma separated list of the sequence lengths to use.
  Default: 1000,10000,100000,500000.

The models will be created in the output directory and should be put into the `data` directory of the PlasClass module to be used. Note that if k-mer and sequence lengths other than the default are used, then these must be specified when calling the `plasclass()` constructor.
